# The Evolution of Human Cancer Gene Duplications across Mammals

**DOI:** 10.1101/2020.03.05.978965

**Authors:** Marc Tollis, Aika K. Schneider-Utaka, Carlo C. Maley

## Abstract

Cancer is caused by genetic alterations that affect cellular fitness, and multicellular organisms have evolved mechanisms to suppress cancer such as cell cycle checkpoints and apoptosis. These pathways may be enhanced by the addition of tumor suppressor gene paralogs or deletion of oncogenes. To provide insights to the evolution of cancer suppression across the mammalian radiation, we estimated copy numbers for 548 human tumor suppressor gene and oncogene homologs in 63 mammalian genome assemblies. The naked mole rat contained the most cancer gene copies, consistent with the extremely low rates of cancer found in this species. We found a positive correlation between a species’ cancer gene copy number and it’s longevity, but not body size, contrary to predictions from Peto’s Paradox. Extremely long-lived mammals also contained more copies of caretaker genes in their genomes, suggesting that the maintenance of genome integrity is an essential form of cancer prevention in long-lived species. We found the strongest association between longevity and copy numbers of genes that are both germline and somatic tumor suppressor genes, suggesting selection has acted to suppress both hereditary and sporadic cancers. We also found a strong relationship between the number of tumor suppressor genes and the number of oncogenes in mammalian genomes, suggesting complex regulatory networks mediate the balance between cell proliferation and checks on tumor progression. This study is the first to investigate cancer gene expansions across the mammalian radiation and provides a springboard for potential human therapies based on evolutionary medicine.

## Introduction

Cancer is a genetic disease arising from the accumulation of mutations that affect cellular survival and division. These mutations, which confer fitness benefits to cellular lineages in the form of greater survival, proliferative potential, and immortality (Hanahan and Weinberg 2011), either arise during somatic evolution within the lifetime of a single individual, or are inherited through the germline and thus responsible for genetic predispositions to cancer (Bodmer and Tomlinson 2010). According to current estimates, between ~1% and ~3.5% of human genes are mutated in cancers (Futreal et al. 2004; Sondka et al. 2018), with ~90% of known cancer-associated mutations occurring somatically, ~20% occurring in the germline, and ~10% occurring as both somatic and germline mutations (Sondka et al. 2018). The types of mutations leading to cancer vary widely, from single point mutations to translocations, but also copy number variations (Stratton et al. 2009) that can include the amplification of oncogenes and/or the deletion of tumor suppressor genes (Nunney 1999; Vogelstein and Kinzler 2004).

While cancer progresses as a consequence of natural selection acting on mutations in cellular lineages, there has also been selection acting at the organismal level to suppress tumor growth, leading to conserved mechanisms such as cell cycle checkpoints and apoptosis (Leroi et al. 2003). Thus, in both the cellular and organismal contexts, cancer is a phenotype that has evolved under multiple levels of selection since the origins of multicellularity (Domazet-Lošo and Tautz 2010; Aktipis et al. 2015). Tumor suppression may be especially challenging in organisms with large bodies and longer lifespans, since cancer is an age- and body size-related disease (Nunney 2018; Seluanov et al. 2018). In humans, longer average leg length results in a higher risk of non-smoking-related cancers (Albanes 1998), and greater than average height has been associated with a higher lifetime risk of melanoma (Lahmann et al. 2016). The probability of getting cancer increases with age in humans (Frank 2007), and greater than 50% of cancers in the US are diagnosed in patients that are 65 and older (White et al. 2014), making advancing age the single biggest risk factor for cancer. Since the number of potentially carcinogenic somatic mutations increases as a function of the number of dividing cells throughout an individual’s lifetime, large multicellular animals with long lifespans theoretically face a higher lifetime risk of cancer than do smaller and shorter lived ones (Peto et al. 1975; Peto 1977). However, across mammals (Abegglen et al. 2015), cancer mortality risk has not been found to be associated with body size or life span: in fact, while the cancer mortality rate in humans is estimated to be between 11% and 25% (Ferlay et al. 2015), it may be as low as 5% in elephants (Abegglen et al. 2015). Thus, the data supports the possibility that larger and long-lived animals actually get less cancer than humans, even though greater size and longevity requires more complex genetic controls over cell growth (Nunney 1999). This suggests that certain animal lineages may have evolved enhanced cancer suppression mechanisms in order to offset tradeoffs associated with large bodies and long lifespans (Roche et al. 2012).

The genetic controls of cancer suppression in nature’s giants has been the focus of much research (Abegglen et al. 2015; Caulin et al. 2015; Sulak et al. 2016; Vazquez et al. 2018; Tollis et al. 2019), as well as in mammals with pronounced longevity such as naked mole rats (*Heterocephalus glaber*) (Seluanov et al. 2009; Tian et al. 2013). Low cancer mortality rates in elephants may be related to the redundancy provided by as many as 20 genomic copies of the tumor suppressor gene TP53 (Abegglen et al. 2015; Caulin et al. 2015; Sulak et al. 2016), which is responsible for apoptosis, senescence, and cell-cycle arrest in the presence of damaged DNA (Kumari et al. 2014). Elephant cells have a higher apoptotic response to DNA damage compared to humans (Abegglen et al. 2015; Sulak et al. 2016), suggesting programmed cell death rather than the repair of damaged DNA is the mechanism of cancer suppression in elephants. Thus, gene duplication can be a major mechanism for the evolution of new traits (Ohta 1989), and redundancies of tumor suppressor genes in large and long-lived animals, such as TP53 in elephants, could effectively require more somatic mutations to trigger malignancies and thus protection from tumors (Nunney 1999). If the evolution of large bodies and long lifespans was accompanied by compensatory and independent anti-cancer adaptations such as tumor suppressor gene duplications, then we should be able to predict lineage-specific tumor suppressor gene expansions in certain lineages. However, to date, the genomic investigations into any link between tumor suppressor gene duplications and the evolution of large bodies or long lifespans have been limited to either a single taxon (Seluanov et al. 2007; Abegglen et al. 2015; Vazquez et al. 2018) or a small number of genomes (Caulin et al. 2015). The recent availability of a large number of diverse mammalian genome assemblies (Koepfli et al. 2015; Lewin et al. 2018) provides an unsourced opportunity to learn about potential genetic mechanisms underlying cancer suppression.

Here, we provide a comparative analysis that examines the relationship between life history traits and the number of cancer gene duplications in a species’ genome. By leveraging the Catalogue of Somatic Mutations in Cancer (COSMIC; Sondka et al. 2018), we estimated the copy numbers of over 500 human cancer gene homologs in 63 complete genomes from across the mammalian radiation. We then performed phylogenetically-informed statistical tests to determine the effects of life history traits such as body mass and longevity on cancer gene copy numbers in mammals. Our study has three goals: 1) to identify mammalian species with the largest numbers of cancer gene paralogs; 2) to determine which life history traits are predictive of cancer gene copy numbers in mammals; and 3) to identify specific mammalian species, or genes, that should be further studied to understand mechanisms of cancer resistance. Combining genomics and trait data, we uncover unique aspects of certain species’ biology and find evidence for multiple, independent mechanisms of cancer suppression across the tree of life, opening up possibilities for future human cancer therapies.

## Results

### Human cancer gene duplications in mammalian genomes

We queried 63 mammalian genome assemblies representing the eutherian superorders Afrotheria, Xenarthra, Euarchontoglires, and Laurasiatheria, and one marsupial (Figure 1A, Supplementary Table 1; Supplementary Materials) for 548 human cancer genes, including 242 tumor suppressor genes and 240 oncogenes (as well as 72 classified as both tumor suppressor and oncogenes) that are causally linked to cancer according to the Cancer Gene Census (CGC) (Sondka et al. 2018) from the COSMIC database (Tate et al. 2019). Within the tumor suppressor genes, 143 genes contained somatic mutations linked to cancer, 35 contained germline mutations, and 43 contained both somatic and germline mutations. To collect homologs for each human cancer gene in mammalian genomes, we performed a protein BLAT search (Kent 2002) using the human peptide, followed by a nucleotide BLAT of the top hit and putative ortholog within the novel genome, and a BLASTX search of each putative paralog to the NCBI human peptide database to ensure a reciprocal match (see Materials and Methods for more details). We then counted the total number of human cancer gene homologs collected for each species, as well as the number of paralogs per species per gene.

Of the 546 successfully validated human cancer gene orthologs, we found that 476 (87.34%) had at least a 1:many relationship (i.e., there are at least two copies of the gene in at least one mammalian genome). We also found that 293 (53.76%) had three copies in at least one mammalian genome, 176 (36.33%) had four copies in at least one mammalian genome, 124 (24.95%) had five copies in at least one mammalian genome, and 45 genes had 10 copies in at least one mammalian genome (8.26%). We did not find an effect of genome assembly length (Phylogenetic Generalized Least Squares Regression or PGLS, Supplementary Figure 1, Supplementary Materials), or assembly completeness (PGLS, Supplementary Figure 2, Supplementary Materials), on estimated cancer gene copy number. mammals, we tested for but found no relationship between estimated cancer gene copy number in a mammalian genome and it’s time to most recent common ancestor (TMRCA) with human as estimated from TimeTree (www.timetree.org) (Kumar et al. 2017).

In order to assess the evolutionary conservation of each human cancer gene ortholog across mammals, we collected 100-way vertebrate phastCons scores (Siepel et al. 2005) for each gene using the UCSC Human Genome Browser (Kent et al. 2002), and validated the presence of each ortholog in the human and either opossum, elephant, or armadillo genome using OrthoDB v9 (Zdobnov et al. 2017). Overall, human cancer genes are not conserved at the sequence level and ranged from 0.001 to 0.9 with a mean of 0.2 (Supplementary Figure 3, Supplementary Materials). We found that only 58 human cancer genes had average phastCons scores above a cutoff of 0.3 which was previously determined to be evolutionarily conserved (Siepel et al. 2005), and that 30 of these (79%) were oncogenes. We found that cancer gene phastCons scores had only a weak effect on their copy number estimates in the human genome (P=0.005, R^2^=0.014) (Supplementary Figure 4, Supplementary Materials). All queried human cancer genes contained orthologs in human and either opossum, elephant, and armadillo. Therefore, while human cancer genes are generally not conserved at the nucleotide level, our results suggest they were present in the common ancestor of eutherians. There were only two human cancer orthologs missing from opossum based on OrthoDB, and we could not detect these genes in the opossum assembly using our BLAT/BLAST search either, suggesting that these orthologs are missing from the opossum assembly.

### Nearly cancer free: Naked mole rats have more paralogous cancer gene copies in their genomes

While we found that many human cancer genes are duplicated in mammalian genomes, the number of predicted paralogous gene copies varied widely per species. For instance, we detected 13 copies of TP53 in the African savannah elephant (*Loxodonta africana*), which is similar to the number of annotated TP53 homologs for this species on Ensembl (Caulin et al. 2015) and what has been reported in other species of elephant (Abegglen et al. 2015; Sulak et al. 2016), while we annotated only a single copy in human. However, the elephant was in the bottom 1/5^th^ for total number of tumor suppressor gene duplications overall (Figure 1B), with an average of 1.65 gene copies per ortholog. The naked mole rat (*Heterocephalus glaber*) contained the most tumor suppressor gene duplications of the queried mammalian genomes, with an average of 2.39 copies per ortholog, followed by two-toed sloth (*Choloepus hoffmanni*; 2.33), human (2.30), and nine-banded armadillo (*Dasypus novemcinctus*; 2.16) (Figure 1B). The naked mole rat also contained the most oncogene duplications (3.40 copies per ortholog), followed by brown rat (*Rattus norvegicus*; 3.36), house mouse (*Mus musculus*; 3.29), and nine-banded armadillo (3.23) (Figure 1C).

**Figure 1.**
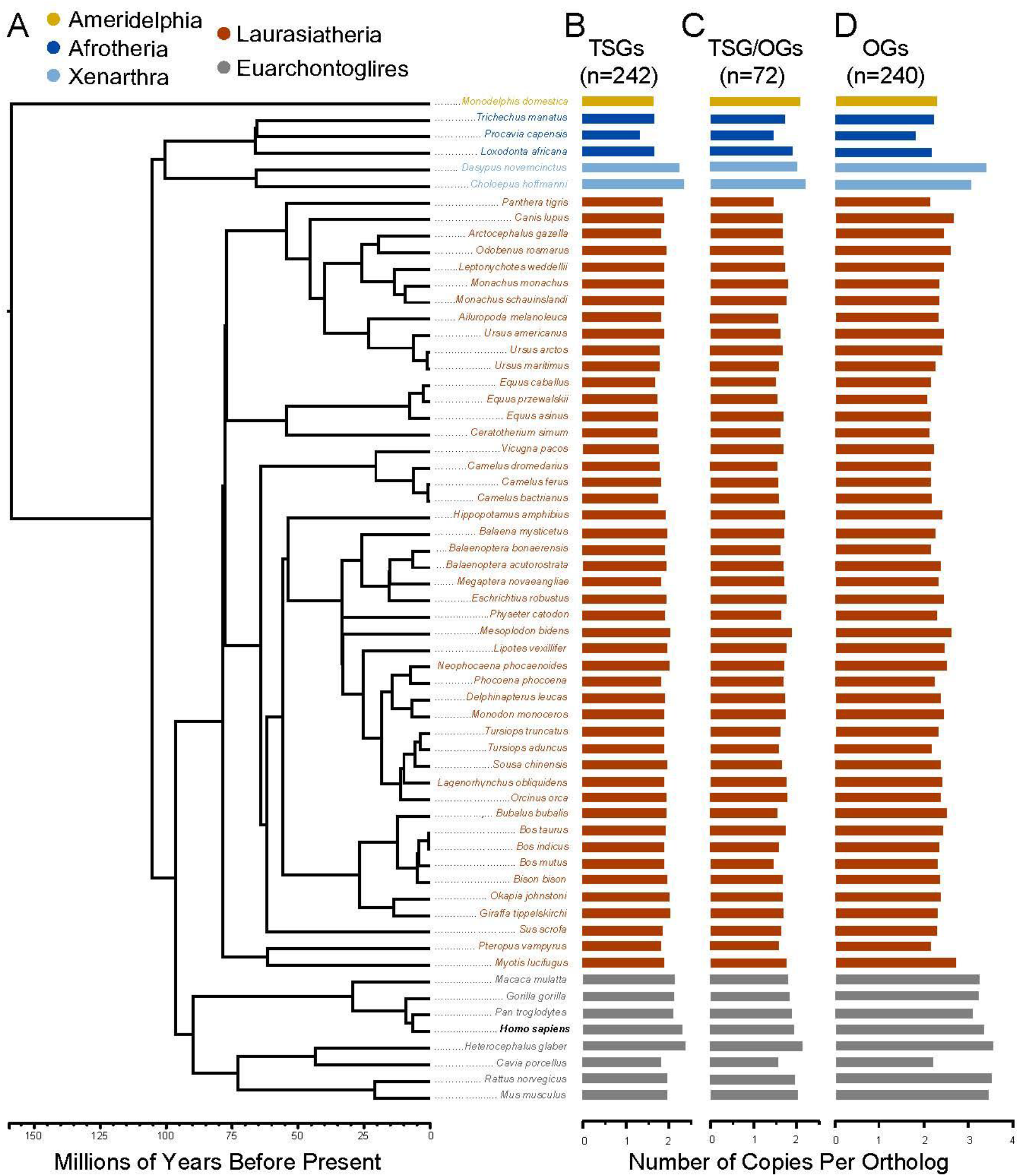
Cancer gene duplications in mammalian genomes. (A) Phylogeny of 63 mammalian genomes used in this study, from Fritz et al. (2009). The tip label for humans (*Homo sapiens*) is in bold. (B) Copy numbers of tumor suppressor genes (TSGs) in each mammalian genome, normalized as the total number of tumor suppressor gene copies divided by the total number of tumor suppressor gene orthologs detected in a genome. (C) Normalized copy numbers for genes that are both tumor suppressor genes and oncogenes (TSG/OGs) in each genome. (D) The normalized copy numbers of oncogenes (OGs). Tumor suppressor gene and oncogene classifications were taken from the Cancer Gene Census (Sondka et al. 2018).

We further divided the tumor suppressor genes by comparing levels of gene duplication in caretaker or gatekeeper genes, according to Caulin et al. 2015. Caretaker genes are responsible for the inhibition of DNA damage as well as the mediation and control of DNA repair, while gatekeeper genes are responsible for the control of cell cycle checkpoints and proliferation (Kinzler and Vogelstein 1997). The mammalian genome with the highest number of caretaker gene duplications was the naked mole rat (2.03), followed by two other euarchontoglires: human (1.61) and brown rat (1.57). The two-toed sloth had the most predicted gatekeeper gene duplications (2.54), followed by naked mole rat (2.42), and human (2.39).

Because natural selection may act differently on somatic mutations compared to germline mutations (Vicens and Posada 2018), we considered an alternative subdivision of tumor suppressor genes depending on whether the cancer associated mutations only occurred somatically, only in the germline, or both. When we compared human cancer gene duplications across the eutherian superorders, we found that the genomes of euarchontoglires (n=8) contained a higher median oncogene and germline tumor suppressor gene duplication, while xenarthran genomes (n=2) contained a higher median duplication of genes that are both tumor suppressors and oncogenes, as well as a higher median duplication when all tumor suppressor genes were classified together (Supplementary Figure 5, Supplementary Materials). In contrast, afrotherian (n=3) and laurasiatherian (n=47) genomes contained lower median duplications of all types of human cancer genes. We found no relationship between cancer gene copy number and TMRCA with humans; therefore, the higher cancer gene copy numbers in rodents are likely due to the fact that euarchontoglires have accumulated more gene expansions compared to other mammals (Demuth et al. 2006) and not related to potential biases stemming from our use of human peptides as query sequences.

### Can’t have one without the other: the balance between tumor suppressor genes and oncogenes

We found a strong correlation between the number of tumor suppressor genes and the number of oncogenes in mammalian genomes (P=2.95e-14, false discovery rate or FDR=3.08e-12, R^2^=0.68), which includes a strong correlation between the number of gatekeeper genes and the number of oncogenes (P=5.81e-14, FDR=3.08e-12, R^2^=0.67), as well as a positive relationship between the number of caretaker genes and the number of oncogenes (P=1.77e-08, FDR=3.13e-08, R^2^=0.47) (Figure 2).

**Figure 2.**
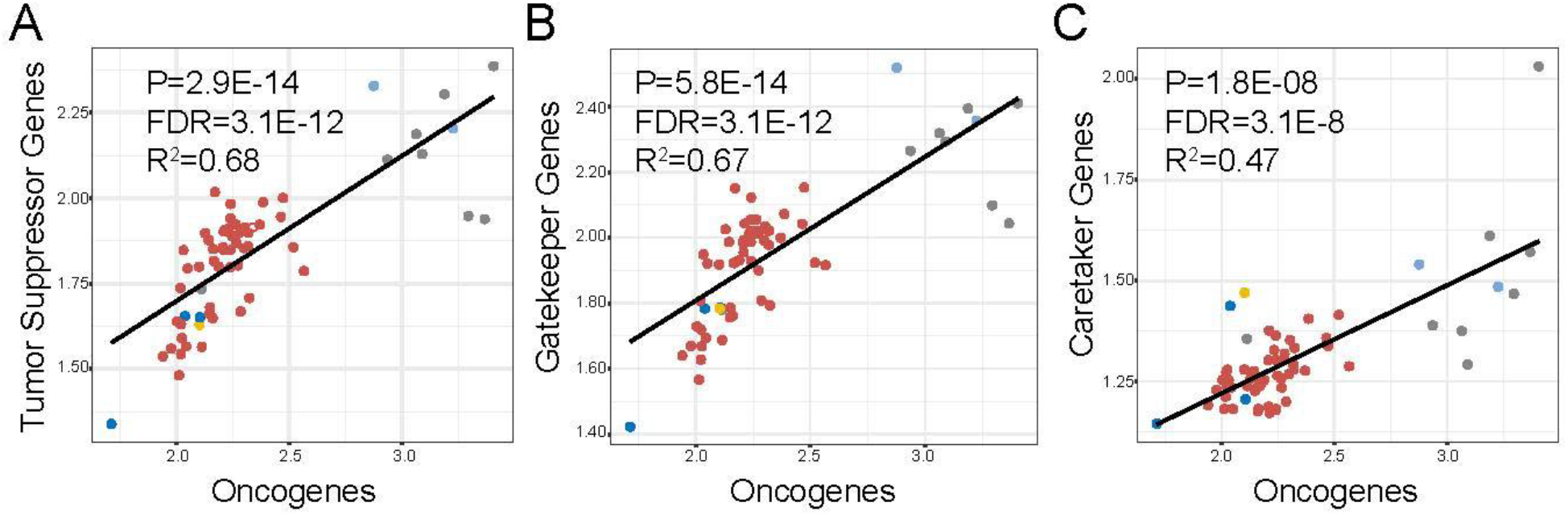
Positive correlations between tumor suppressor gene and oncogene copy numbers in 63 mammalian genomes. (A) Phylogenetic least squares regression (PGLS) between copy numbers of all tumor suppressor genes and copy numbers of oncogenes in mammalian genomes. (B) PGLS between copy numbers of caretaker genes and copy numbers of oncogenes. (C) PGLS between copy numbers of gatekeeper genes and copy numbers of oncogenes. Colors represent mammalian clades following Figure 1A. All y-axes are given in terms of normalized cancer gene copy number (the total number of cancer gene homologs divided by the number of found cancer gene orthologs). FDR=false discovery rate.

### Longevity matters more than just size

Mammalian megafauna such as elephants and whales are made of, respectively, 100X and 1000X more cells than smaller human-sized mammals. Such a large number of cells may confer a greater risk of cancer-causing mutations during somatic evolution (Caulin et al. 2015). Therefore, large-bodied mammals may have evolved compensatory adaptations for extra cancer defenses to ensure survival through reproductive age (Peto 1977, Caulin and Maley 2011, Abegglen et al. 2015). We hypothesized that a potential genetic mechanism for cancer suppression in large mammals could be a redundancy of tumor suppressor genes that would effectively require more “hits” to trigger carcinogenesis (Nunney 1999, Caulin and Maley 2011). Conversely, we expected to find a paucity of oncogenes in the genomes of larger mammals relative the genomes of smaller mammals. To test this, we collected body mass (in grams) and lifespan (in years) data for the 63 mammalian species in this study (Figure 3).

**Figure 3.**
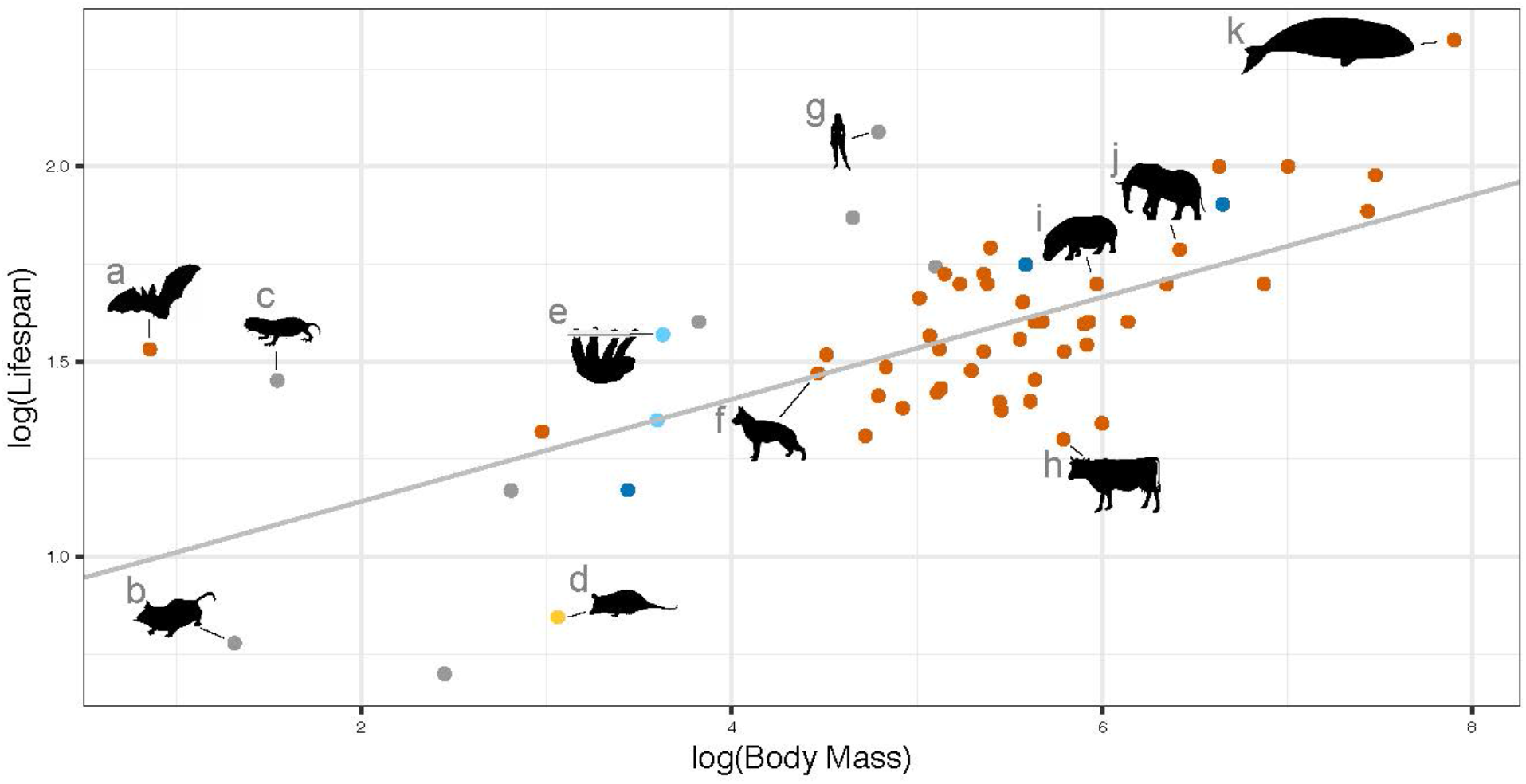
Relationship between body mass (g) and lifespan data (years) for 63 mammals. The grey line represents a linear regression between log_10_ body mass and log_10_ maximum lifespan (slope = 0.13, intercept = 0.88). Colors represent mammalian clades following Figure 1A. Silhouettes from PhyloPic.org represent key lineages (animal figures not drawn to scale): (a) little brown bat (*Myotis lucifugus*), (b) house mouse (*Mus musculus*), (c) naked mole rat (*Heterocephalus glaber*), (d) grey opposum (*Monodelphis domestica*, by Sarah Werning https://creativecommons.org/licenses/by/3.0/), (e) two-toed sloth (*Choleopus hoffmanni*), (f) dog (*Canis lupus familiaris*), (g) human (*Homo sapiens*), (h) cow (*Bos taurus*), (i) hippopotamus (*Hippopotamus amphibius*), (j) African bush elephant (*Loxodonta africana*, by Jan A. Venter, Herbert H. T. Prins, David A. Balfour & Rob Slotow, vectorized by T. Michael Keesey, https://creativecommons.org/licenses/by/3.0/), (k) bowhead whale (*Balaena mysticetus*, by Chris Huh https://creativecommons.org/licenses/by-sa/3.0/).

We found no relationship between the number of tumor suppressor gene duplications and body mass across mammals when we considered all tumor suppressor genes together (Figure 4A); however, we did find that, surprisingly, copy numbers of tumor suppressor genes which are mutated in both germline and soma are negatively correlated with body mass (P=0.002, FDR=0.01, R^2^= 0.18, Figure 4B). We also found a negative relationship between body size and oncogene copy numbers (P=0.04, R^2^=0.08, Figure 4C). However, these results did not pass significance criteria after applying the false discovery rate (FDR) to account for multiple testing (see Materials and Methods), or when taking phylogeny into account using a PGLS regression. Therefore, larger mammals did have fewer oncogenes in their genomes than smaller mammals, albeit not to a statistically significant degree. We found no evidence that larger mammals contain more tumor suppressor genes.

As cancer is a disease of not just body size but also longevity, we tested for a relationship between human cancer gene copy numbers and longevity in 63 mammals (see Materials and methods). Briefly, we collected maximum body mass and lifespan data for 2,549 mammals in order to calculate longevity quotients (LQ) for each species. LQ gives an indication of how long a species’ lifespan is compared to other species of similar size, where LQ=observed longevity/expected longevity (Austad and Fischer 1991). We calculated the expected longevity for each species by fitting a linear regression to log_10_(maximum longevity) and log_10_(body mass) using non-flying eutherian mammals, following Austad and Fischer (1991) and Foley et al. (2018), obtaining a slope of 0.27 and an intercept of 0.14 (R^2^=0.51) (Figure 4D). We were unable to obtain lifespan data for Sowerby’s beaked whale (*Mesoplodon bidens*); therefore, we did not calculate LQ for this species.

**Figure 4.**
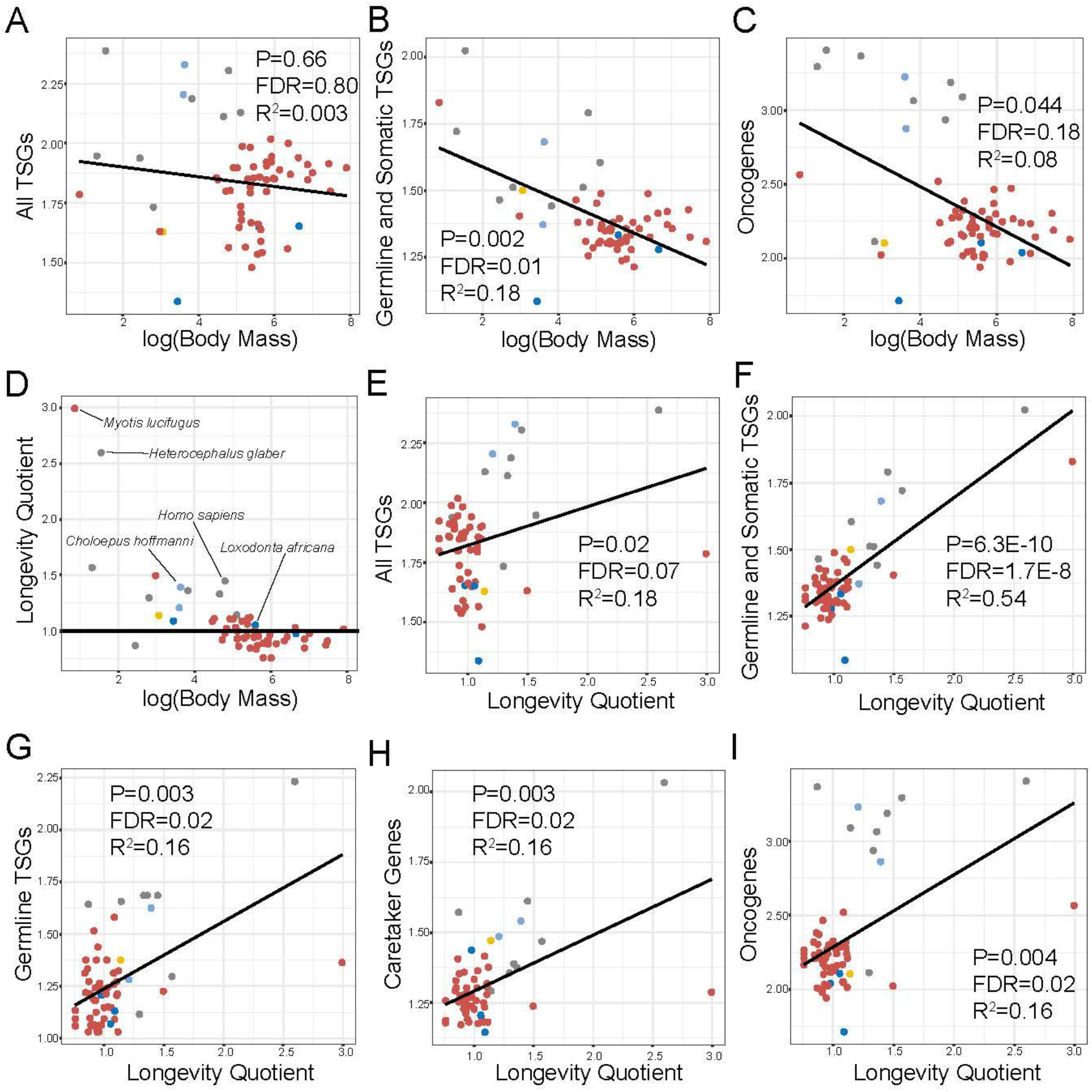
Cancer gene duplications are positively associated with longevity, and not body mass, in mammalian genomes. (A) There is no association between tumor suppressor gene (TSG) duplications and body mass (given in log_10_) when we grouped all TSG types together. (B) The only subset of TSGs whose copy numbers in mammalian genomes are significantly associated with body mass are those which are mutated in both human germline and soma. This relationship is slightly negative and is of moderate significance (FDR or false discovery rate=0.01). (C) Larger mammals have fewer oncogene duplications; however, this relationship in non-significant when taking into account both phylogeny and multiple testing. (D) Longevity quotients (LQ) versus body mass for 62 mammals included in this study. Key species relevant to this study are labeled. Black line indicates LQ=1.0, where the observed longevity is equal to the expected longevity. (E) There is a positive correlation between TSG duplications and LQ when we grouped all TSG types together; however this relationship is non-significant when taking into account multiple testing (FDR=0.07). (F) The strongest correlation (R^2^=0.54) was found between duplications in TSGs mutated in both human germline and soma and longevity. LQ is also correlated with duplications in TSGs with only germline mutations (G), caretaker genes (H), and oncogenes (I). Colors represent mammalian clades following Figure 1A. All y-axes are given in terms of normalized cancer gene copy number (the total number of cancer gene homologs divided by the number of found cancer gene orthologs).

Across mammals, we found a positive relationship between copy numbers of all tumor suppressor genes and LQ (Figure 4E), although this result did not pass significance criteria after correcting for multiple testing. We found the strongest (and most statistically significant) positive relationship between LQ and copy numbers of tumor suppressor genes mutated in both germline and soma (P=6.26e-10, FDR=1.7e-8, R^2^=0.54) (Figure 4F). We also found significant relationships between LQ and copy numbers of germline tumor suppressor genes (P=0.003, FDR=0.02, R^2^=0.16, Figure 4G), caretaker genes (P=0.003, FDR=0.02, R^2^=0.16, Figure 4H), and oncogenes (P=0.004, FDR=0.02, R^2^=0.17, Figure 4I). All of these tests passed the criteria for significance after correcting for multiple testing. We found no effect of basal metabolic rate, diet, placental invasiveness or biome on cancer gene copy number (Supplementary Figure 6, Supplementary Materials).

In order to determine whether these patterns were unique to cancer genes, we also estimated copy numbers for curated sets of 87 housekeeping genes and 214 nervous system genes from (Dorus et al. 2004) (Supplementary Figure 7 and Supplementary Figure 8, Supplementary Materials). We found no relationship between any of the life history variables and housekeeping gene copy number. We found a weak effect of body mass (P=0.02, R^2^=0.11) and LQ (P=0.02, R^2^=0.10) on nervous system gene copy number; however these results did not pass significance criteria after correction for multiple testing. Therefore, according to our results longevity had a striking and unique effect on the number of cancer gene copies in a mammalian genome, and the strongest predictor of cancer gene copy number in a mammalian genome was its lifespan relative to other mammals of similar size.

## Discussion

The evolutionary relatedness of other mammals to humans has led to the realization that patterns of tumor suppressor gene duplication in mammalian genomes may shed light on cancer resistance mechanisms and lead to novel therapies (Caulin and Maley 2011; Abegglen et al. 2015; Caulin et al. 2015; Keane et al. 2015; Tollis et al. 2017; Vazquez et al. 2018). However, to date, the few studies that have queried mammalian genomes for tumor suppressor gene duplications have mostly focused on few taxa (Caulin et al. 2015; Keane et al. 2015) or a small number of genes (Abegglen et al. 2015; Sulak et al. 2016; Vasquez et al. 2018). Here, we have provided the most comprehensive survey of cancer gene duplications across the mammalian radiation to date, incorporating 548 known human cancer genes and 63 mammalian genomes. Our results show that ~87% of human cancer genes are duplicated in at least one mammalian genome, and that there are many lineage-specific cancer gene expansions.

A caveat of this study is that it is impossible to identify true biological gene deletions in draft genomes which may harbor assembly errors and varying levels of incompleteness. Nonetheless, we found no relationship between assembly length or predicted assembly completeness and cancer gene copy number. Another potential problem in our analysis is the possibility of an ascertainment bias that stems from using human peptides as query sequences; evolutionarily distant mammalian genomes could contain undetected homologs due to high levels of sequence divergence. Several results suggest this was not a problem for our study: (1) we found no relationship between the TMRCA with human and cancer gene copy number in our mammalian genome assemblies; (2) all queried human cancer genes shared orthologous matches in every tested mammal, with only two exceptions in the marsupial, suggesting widespread orthology; and (3) we found only a weak relationship between evolutionary sequence conservation and cancer gene copy number. All these results suggest that our ability to detect cancer gene copies in a given mammalian genome did not rely on evolutionary distance from humans. Other caveats are the lack of experimental evidence that the human cancer genes in this study share the same function in other species, and that there may be species-specific pathways for tumor suppression in addition to those causally linked to human cancers. Thus, our results must be taken with some caution until these are experimentally verified.

Our finding that the naked mole rat has the most cancer gene duplications is consistent with previous studies suggesting that naked mole rats have very strong cancer defenses and anti-aging adaptations (Buffenstein 2005; Buffenstein 2008; Seluanov et al. 2009; Tian et al. 2013), and that until recently (Delaney et al. 2016) there have been no reported cases of cancer in the species. In fact, we found that rodents such as naked mole rat, brown rat, and house mouse contained more copies of every category of cancer gene than other mammals, except for the two-toed sloth which had the most gatekeepers, somatic tumor suppressor genes, and genes that are both tumor suppressors and oncogenes. Rodents contain a diversity of lifespan phenotypes that range from 4 years in mice to over 30 years in naked mole rats. Surprisingly, it was found that the repression of telomerase activity, which is a well studied anticancer mechanism, was related to body mass rather than lifespan in rodents (Seluanov et al. 2007), suggesting that larger species must mitigate the risk of tumorigenesis that stems from cell proliferation. While we found support for a positive relationship between tumor suppressor gene copy number and body mass and lifespan in euarchontoglires (P=0.02, R^2^=0.69) (Supplementary Figure 5, Supplementary Materials), these tests did not pass significance criteria after correction for multiple testing. Still, the naked mole rat had both the highest longevity quotient and the most cancer gene duplications, meaning our results make it difficult to reject a correlation between body mass, lifespan, and anticancer adaptations in rodents.

The two-toed sloth and nine-banded armadillo also contained a comparatively large number of cancer gene duplications. These species are xenarthran mammals which diverged from other eutherians during the early Cenozoic (Meredith et al. 2011), making them only very distantly related to rodents and primates. To our knowledge, there has not been any cancer reported in sloths; however, sloth fur carries an anti-cancer fungus (Higginbotham et al. 2014). There have been reports of cancer in armadillos (Pence et al. 1983; Lee et al. 2015). Neither of these species are particularly large-bodied, nor do they exhibit remarkable longevity according to our estimates. Therefore, it is not clear why these xenarthrans would have special anticancer defenses. Among mammals, xenarthrans are thought to have the lowest basal metabolic rate (McNab 1985; Pauli et al. 2016), which may be a potential anti-cancer mechanism for cancer-resistant species (Caulin and Maley 2011) due to reduced oxidative damage to DNA which could otherwise trigger tumor formation (Adelman et al. 1988; Xie et al. 2004). However, we found no relationship between basal metabolic rate and cancer gene copy number. We suggest that sloths and armadillos should be the focus of future studies investigating cancer resistance.

Bats (order Chiroptera), on the other hand, have both high metabolic rates and exceptional longevity (Munshi-South and Wilkinson 2010) (Figure 4D). We found that the little brown bat (*Myotis lucifugus*) was an outlier compared to other laurasiatherians with high cancer gene copy numbers, in particular for tumor suppressor genes with both germline and somatic mutations (1.83 copies per ortholog versus 1.42 average for laurasiatherians). Recent comparative studies have suggested that bats in the long-lived genus *Myotis* do not display telomere shortening with age (Foley et al. 2018), showed evidence of positive selection in genes that control DNA damage (Foley et al. 2018), and have unique gene expression patterns in pathways controlling DNA repair and tumor suppression that increase with age (Huang et al. 2019). Similarly, a previous study found accelerated protein evolution in DNA damage and response pathways that was unique to long-lived mammals (Li and de Magalhães 2013). Meanwhile, we found that copy numbers of caretaker genes were significantly correlated with longevity across mammals. Altogether, these results are consistent with other studies (MacRae et al. 2015) and suggest that the maintenance of genome integrity may be an essential form of cancer prevention in long-lived species.

Copy numbers of cancer genes bearing both germline and somatic mutations according to COSMIC yielded the strongest positive correlation with longevity, followed by germline-only-mutated cancer genes. This suggests that pathways associated with hereditary cancers, which are caused by germline mutations, may face different constraints than those associated with sporadic cancers. Vicens and Posada (2018) analyzed ratios of the proportion of nonsynonymous substitutions per site to the proportion of synonymous substitutions per site in human cancer genes across mammals and found evidence for relaxed selective constraint in cancer genes with germline mutations relative to those with somatic mutations. Thus, there may be stronger purifying selection acting on pathways associated with highly deleterious sporadic cancers, which would reduce the fixation rate of copy number variants related to those pathways. Still, hereditary cancers are likely strongly selected against in long-lived species due to the fact that in general cancer is a late-onset disease. Our results suggest that additional copies of genes with both germline and somatic mutations linked to cancer may provide an extra safeguard against both hereditary and sporadic cancers in long-lived species.

Interestingly, we found that copy numbers of both tumor suppressor genes and oncogenes were positively correlated with longevity across mammals. At the same time, the number of tumor suppressor genes in a genome was strongly correlated with the number of oncogenes. Caulin et al. (2015) found a positive correlation between the number of oncogenes and the number of gatekeepers, while we found the positive correlation also holds for caretaker genes. It is possible that Caulin at al. (2015) may not have had the statistical power to detect this relationship. Their study lacked several mammalian genomes included here, such as the naked mole rat which, according to our results, contains the most caretaker genes. It seems logical that long-lived species would have more copies of tumor suppressor genes, which control the cell cycle and programmed cell death, thus inhibiting tumor growth (Vogelstein and Kinzler 2004). Meanwhile, oncogenes are genes that, when amplified, activate and promote cell growth and proliferation. It seems paradoxical, therefore, that there would also be oncogene expansions in long-lived species. However, longer-lived species may have more complex regulatory networks that mediate the balance between cellular proliferative potential and necessary checks on tumor progression. In the human and mouse genomes, oncogenes without a tumor suppressor gene at a nearby genomic location were found more likely to become amplified (Wu et al. 2017). Oncogenes and tumor suppressor genes were found to be among the evolutionarily oldest sets of genes in eukaryotes, associated with the timing of the origins of both multicellular and vertebrate life (Makashov et al. 2019); this is consistent with our observation that oncogenes are highly conserved at the sequence level. The co-amplification of oncogene and tumor suppressor gene families may be the result of a positive feedback system in which the addition of tumor suppressor genes is an inhibitory response to the expansion of oncogenes, or vice-versa.

In conclusion, our study has shed light on many aspects of the evolution of cancer suppression in mammals. The naked mole rat genome contains more cancer gene copies than all of the other mammals we investigated, even after correcting for phylogeny and ascertainment biases, which is consistent with the very low cancer rates found in this species. We also found that while euarchontoglires (the mammalian superorder that include primates and rodents) have the most cancer gene copies, there have also been significant cancer gene duplications in xenarthrans (sloths and armadillos), and that laurasiatherians contain comparatively much fewer cancer gene copies with the little brown bat as an outlier. We suggest that these species should be the focus of future functional assays that investigate potential cancer defense mechanisms. We found that contrary to predictions stemming from Peto’s Paradox, body size does not explain much of the variance in mammalian cancer gene copy number, while longevity does. Tumor suppressor genes that are mutated in both the germline and the soma had significantly more duplications in the genomes of mammals with pronounced longevity, including the little brown bat and the naked mole rat, suggesting extra cancer defenses against both heritable and spontaneous cancers in long-lived species. Our study adds to the growing body of evidence that adaptations for cancer defense mechanisms have occurred across the mammalian tree of life, and that mammals with exceptional longevity relative to their body size may hold genetic keys to cancer prevention.

## Materials and methods

### Collection of orthologs

A list of human cancer genes, including tumor suppressor genes and oncogenes, was downloaded from COSMIC (last accessed December 2018). For each cancer gene, we retrieved (1) gene symbols, (2) gene names, and (3) whether cancer-related mutations were germline, somatic, or both. Each tumor suppressor gene was given the classification of being a gatekeeper gene or a caretaker gene according to the 59 gatekeepers and caretakers reported in Caulin et al. 2015; we classified the remaining tumor suppressor genes as either gatekeepers or caretakers using the descriptions for each gene available from GeneCards (Stelzer et al. 2016) (Supplementary Materials). We then gathered human (GRCh38.p13) protein sequences for each human cancer gene ortholog from Ensembl v98 using BioMart (Kinsella et al. 2011). The complete genome assemblies of 63 mammals were collected from NCBI (O’Leary et al. 2016) and The Bowhead Whale Genome Resource (Keane et al. 2015). Each genome assembly was assessed for completeness by calculating the proportion of complete universal single copy orthologs from mammalia_odb10 with BUSCO v4.0 (Simão et al. 2015).

To collect human cancer gene orthologs in mammalian genomes, we used the human peptide in a protein BLAT search with minscore=55 and minidentity=60. To collect cancer gene paralogs, we used the DNA sequence of the top scoring hit from the protein BLAT in a second BLAT search of the same genome. We then used each putative paralogous hit at least 33.3% of the length of the coding sequence of the human ortholog as a query in a BLASTX (Boratyn et al. 2012) of the RefSeq human protein database (taxid:9606), keeping only copies that returned a top hit of the original human peptide with ≥65% protein identity. This approach was more conservative than using the BLAT parameters on the UCSC Genome Browser web server; however, using web BLAT settings resulted in a large number of false positives (i.e. the reciprocal BLASTX returned the wrong human ortholog). Whenever the first search resulted in either missing homologs or a large number of false positives, we reran each analysis with either more relaxed BLAT search parameters and/or a slightly more conservative protein identity cutoff for BLASTX (≥70%).

Despite relaxing the parameters, two cancer genes (PRDM1 and QKI) consistently returned zero copies. We therefore performed the bioinformatic search steps for these genes manually using web BLAT on 33 mammalian genomes hosted on the UCSC Genome Browser, and ultimately obtained verified homologs for both genes. To account for potential missing orthologs due to incomplete genome sequencing and assembly, we normalized the cancer gene copy number estimates for each mammalian genome to equal the total number of cancer gene homologs divided by the number of found cancer gene orthologs. To estimate evolutionary conservation in human cancer genes, we averaged the per-site phastCons scores (phastCons100way, Siepel et al. 2005) for the Ensembl whole gene annotations of every human cancer gene ortholog using the UCSC Table Browser (Karolchik et al. 2004).

Dorus et al. (2004) provided a list of 94 housekeeping genes, seven of which were also found in COSMIC and were thus removed from analysis, as well as 214 nervous system genes. The human peptides for these genes were used as queries in a bioinformatic search as described above using the 63 mammalian genomes.

### Collection of trait data

We collected maximum body mass and lifespan data for the 63 mammals whose genomes were analyzed using the Amniote Life History Database (Myhrvold et al. 2015). Additionally, we collected mean basal metabolic rate (BMR, ml O2/hr) for 25 of these species, 19 of which were previously reported (Wilson Sayres et al. 2011). We were able to obtain BMR from six more of our studied species from additional sources (Gumal and Irwan 1998; Jones et al. 2009; Sieg et al. 2009). We collected additional data for each species including biome (tropical or non-tropical), habitat (aquatic or terrestrial) and diet (carnivore, omnivore or herbivore) from Animal Diversity Web (Meyers et al. 2020). We obtained placentation data (hemochorial, endotheliochorial or epitheliochorial) for 31 mammals from Comparative Placentation (http://placentation.ucsd.edu/homefs.html) and Mossman (1987). The life history data as well as the cancer gene copy number estimates for all 63 mammals is in Appendix A.

### Longevity quotient calculations

To increase the number of data points compared to previous studies of LQ (Austad and Fischer 1991; Foley et al. 2018), we accessed the Amniote Life History Database and collected maximum body mass and lifespan data for 2,549 mammalian species including 2,060 non-flying eutherians, 260 chiropterans, and 299 marsupials. We calculated the expected longevity of each species by fitting a linear regression to log_10_(maximum longevity) and log_10_(body mass) using the non-flying eutherian mammal data as in Austad and Fischer (1991) and Foley et al. (2018). Calculations were performed in R v3.5.1 (R Core Team 2018), and we accounted for phylogenetic dependence in the data using *caper* v1.0.1 (Orme et al. 2018) and a mammalian supertree phylogeny (Fritz et al. 2009).

### Phylogenetically-informed regressions of cancer gene copy numbers and trait data

We tested the relationships between normalized human cancer gene copy number (for total tumor suppressor genes, germline tumor suppressor genes, somatic tumor suppressor genes, gatekeepers, caretakers, oncogenes, and genes that are both tumor suppressors and oncogenes) and the trait data. In total, we completed 106 PGLS regressions using *caper* v1.0.1 and the mammalian supertree phylogeny, available in Appendix B. We applied a false-discovery rate where q=0.05 in order to account for multiple testing (Mcdonald 2014).

## Data availability

All sequence data from this study is made publicly available at the Harvard (https://doi.org/10.7910/DVN/3CVMQI). The automated bioinformatic pipeline is made available at https://github.com/bhanratt/orthologcollector.

## Acknowledgements

This work was supported in part by NIH grants U54 CA217376, U2C CA233254, P01 CA91955, R01 CA149566, R01 CA170595, R01 CA185138 and R01 CA140657 as well as CDMRP Breast Cancer Research Program Award BC132057 and the Arizona Biomedical Research Commission grant ADHS18-198847 (CCM). We would like to thank Brian Hanratty from the Bioinformatics Core at Arizona State University. We also extend gratitude towards Dr. Melissa Wilson (Arizona State University) for valuable comments during early manuscript preparations.

