## Supplementary Materials for "The Evolution of Human Cancer Gene Duplications across Mammals"

**Supplementary Figure 1. Estimates of tumor suppressor gene and oncogene copy number variants are not related to genome assembly length.** Tumor suppressor gene and oncogene duplications are given as the number of total genes detected in the genome divided by the number of orthologous genes detected in the genome.

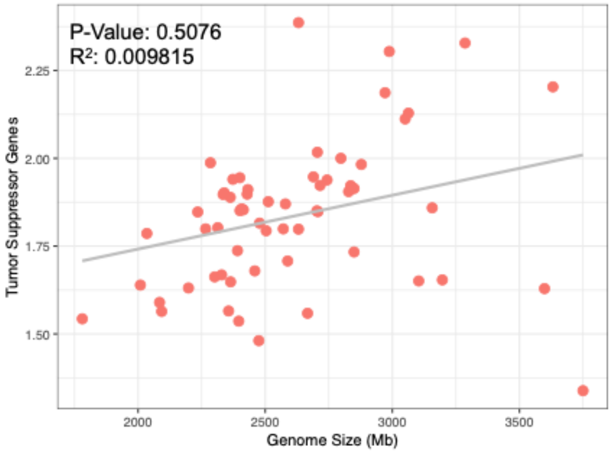


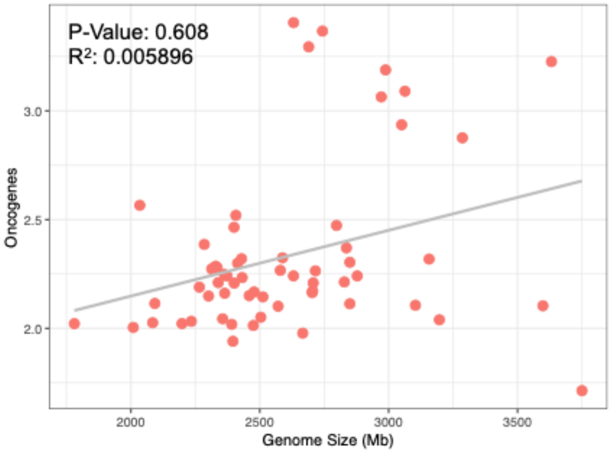

**Supplementary Figure 2. Estimates of tumor suppressor gene and oncogene copy number variants are not related to the number high-confidence orthologs found in a genome assembly.** Tumor suppressor gene and oncogene duplications are given as the number of total genes detected in the genome divided by the number of orthologous genes detected in the genome. BUSCO indicates the percentage of 9,226 mammalian single copy orthologs that were complete in each genome assembly.

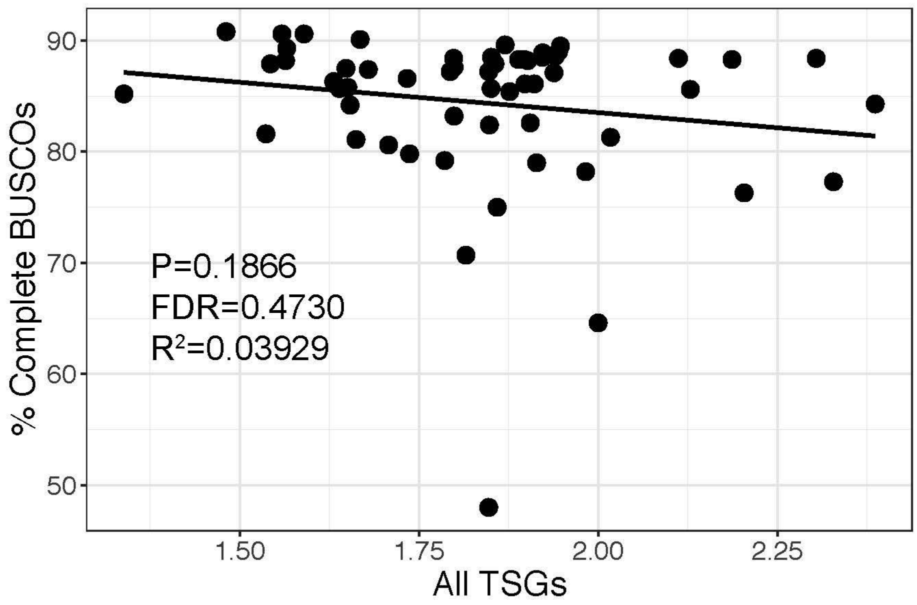


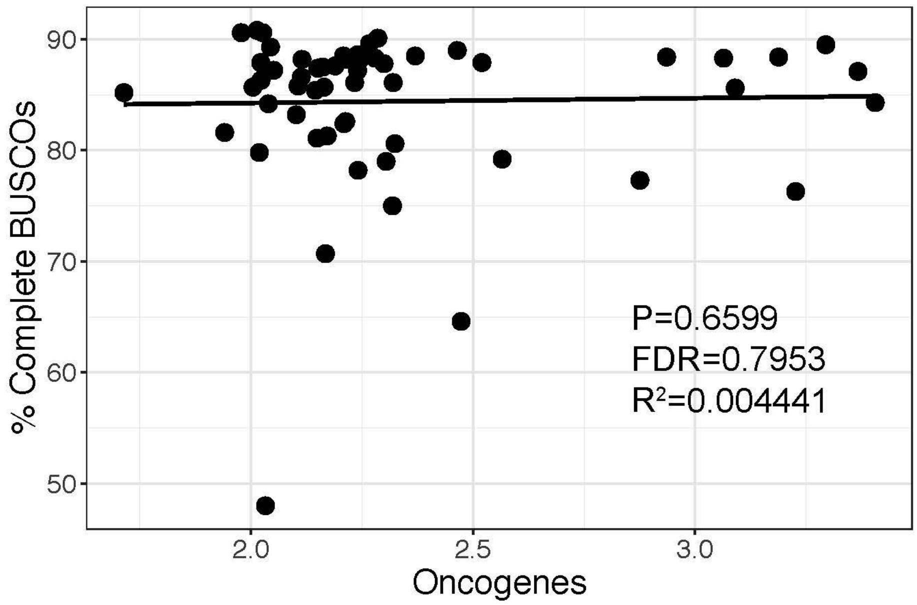

**Supplementary Figure 3. Distribution of average phastCons (vertebrate 100-way) scores for human cancer genes.**

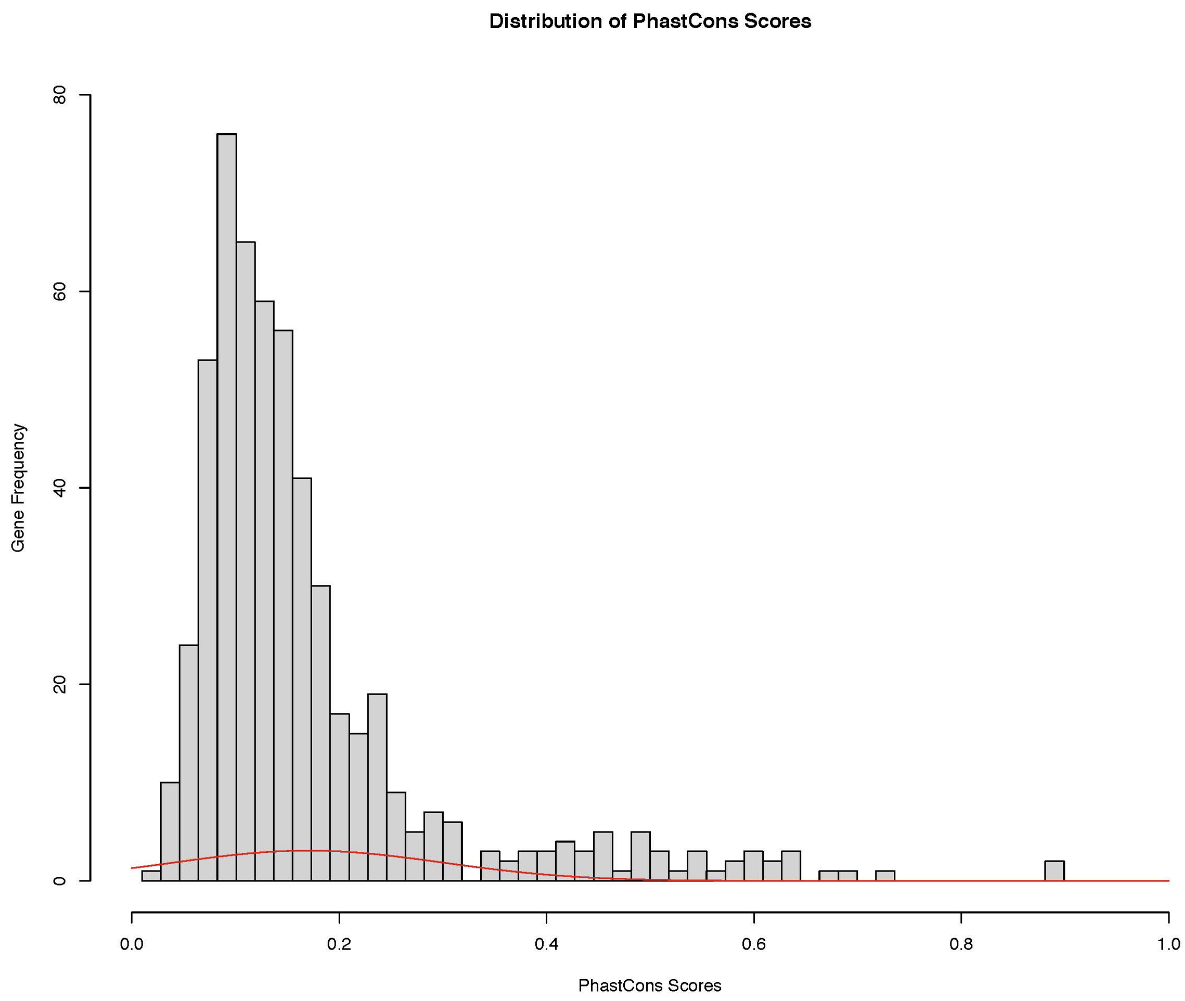

**Supplementary Figure 4. Cancer gene phastCons scores have a weak effect on their copy number estimates in the human genome.**

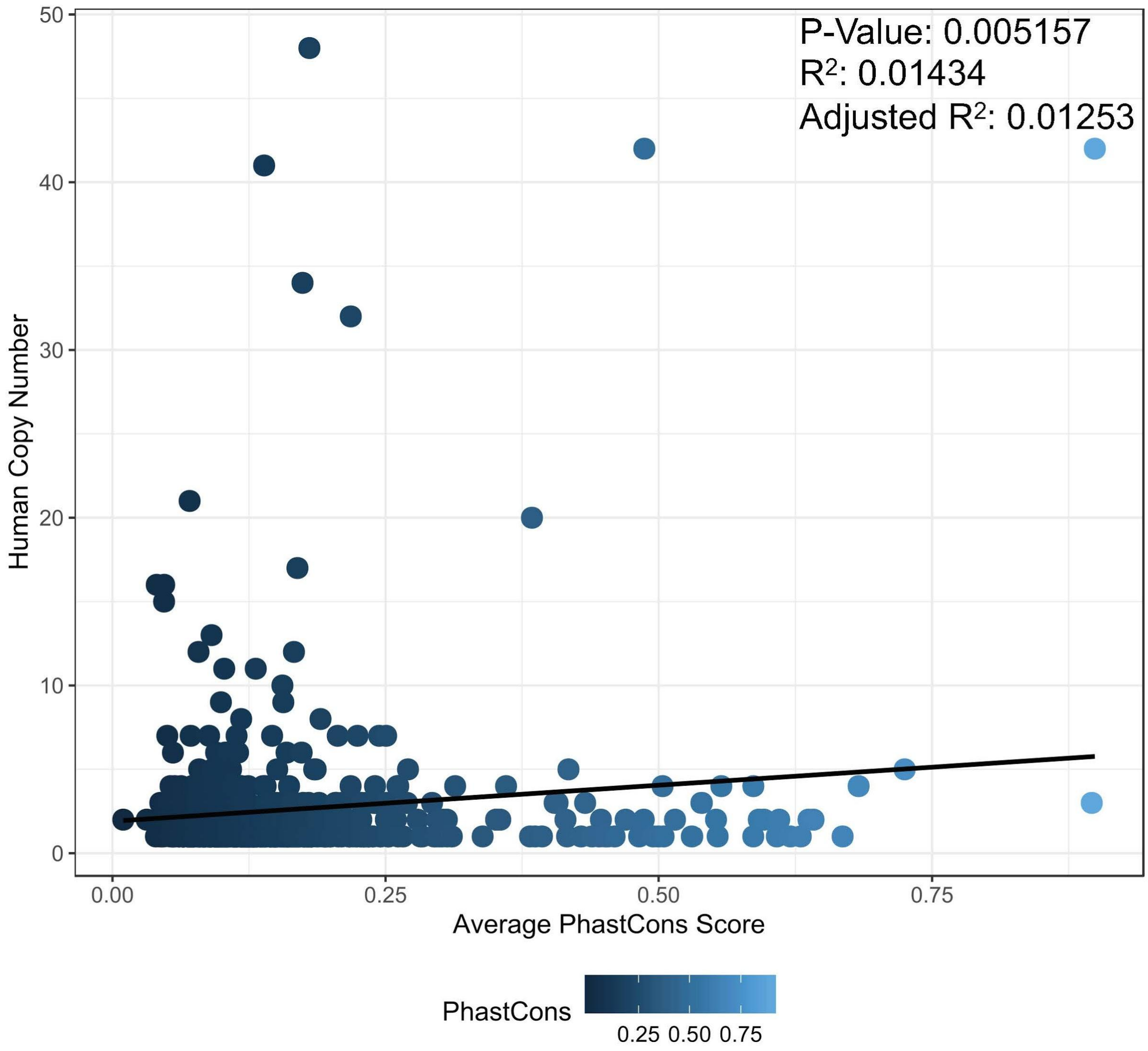

**Supplementary Figure 5. Cancer gene duplications differ across mammalian clades.**

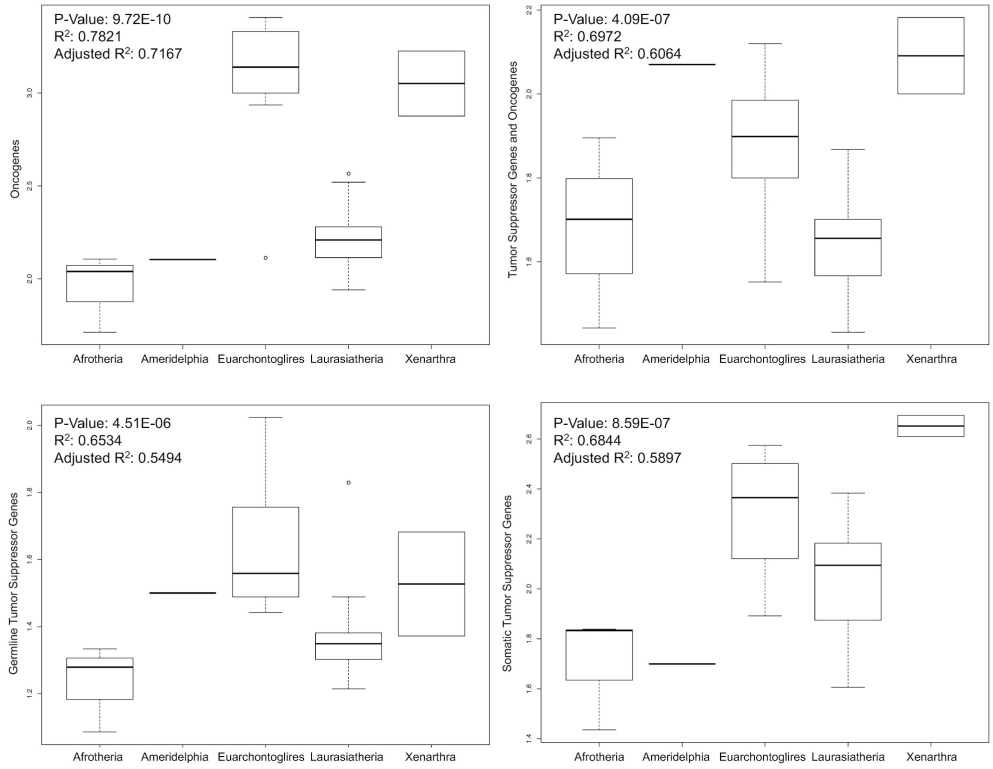

**Supplementary Figure 6. Human cancer gene copy numbers are not correlated with basal metabolic rate (mL $O_2$/hr) across mammals.**

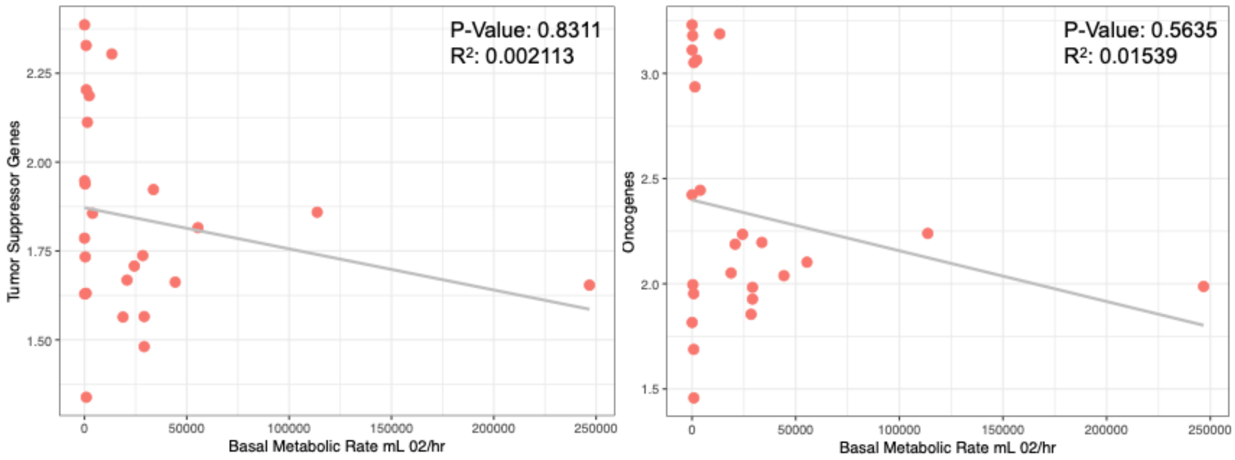

**Supplementary Figure 7. Copy numbers of housekeeping genes are not correlated with body mass or longevity in 63 mammalian genomes.**

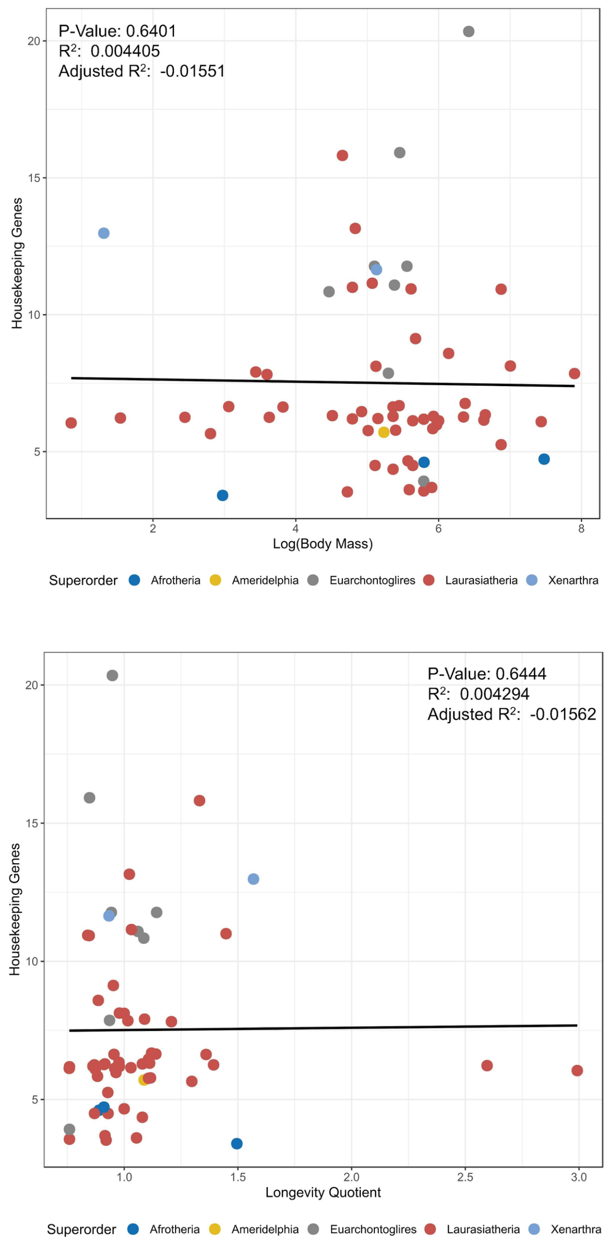

**Supplementary Figure 8. Nervous system gene copy numbers are weakly correlated with body mass and longevity in 63 mammals.** This correlation is non-significant when accounting for multiple testing.

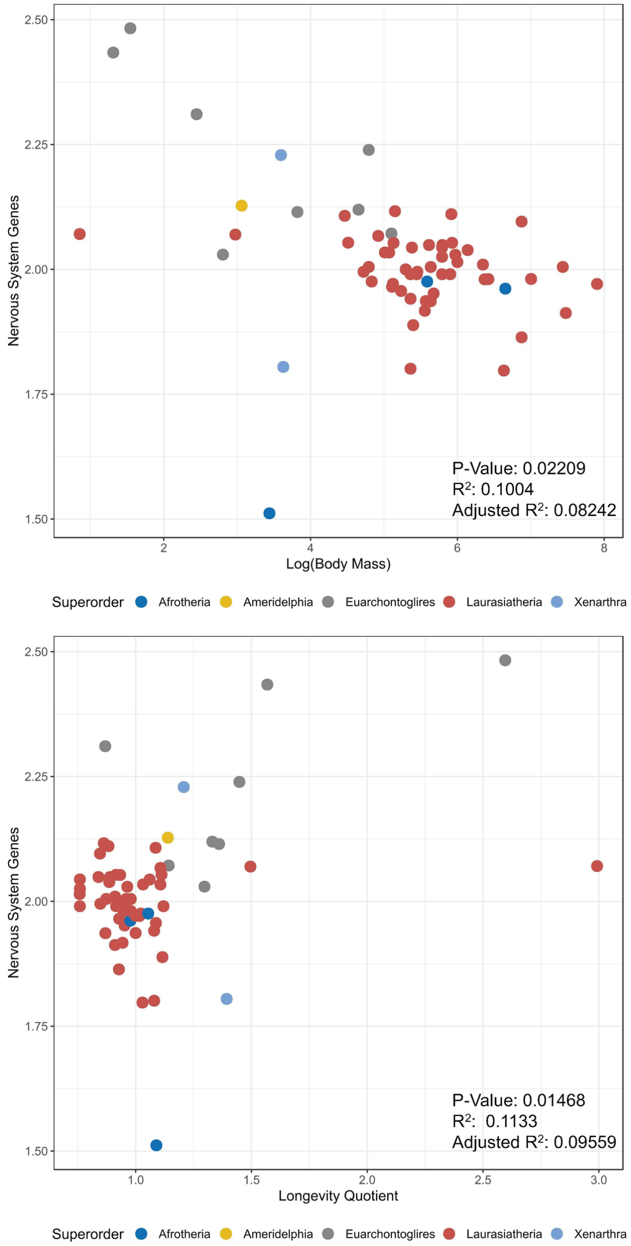

### Supplementary Table 1. Genome assemblies accessed for this study.

| Species Name | NCBI Assembly ID | Sex | Genome Length | Genome Coverage | Sequencing Method | Scaffolds or Chromosomes |
|---|---|---|---|---|---|---|
| *Vicugna pacos* Alpaca | Vi_pacos_V1.0 | Female | 2092.95 Mb | 72.5x | Illumina HiSeq2000 | Scaffolds |
| *Bison bison* American Bison | Bison_UMD1.0 | Male | 2828.03 Mb | 60x | 454; Illumina HiSeq | Scaffolds |
| *Arctocephalus gazella* Antarctic Fur Seal | ArcGazv1.4 | Female | 2313.49 | 200x | Illumina HiSeq | Scaffolds |
| *Balaenoptera bonaerensis* Antarctic Minke Whale | ASM97880v1 | N/A | 2234.64 | 60x | Illumina HiSeq2000 | Scaffolds |
| *Camelus dromedarius* Arabian Camel | PRJNA234474_Ca_dromedarius_V1.0 | Male | 2084.54 | 46.43x | Illumina HiSeq2000 | Scaffolds |
| *Dasypus novemcinctus* Nine-Banded Armadillo | Dasnov3.0 | Female | 3631.52 | 6x | Sanger | Scaffolds |
| *Camelus bactrianus* Bactrian Camel | Ca_bactrianus_MBC_1.0 | Female | 1780.72 | 79.2x | Illumina HiSeq2000 | Scaffolds |
| *Delphinapterus leucas* Beluga Whale | ASM228892v3 | Female | 2362.78 | 117x | Illumina HiSeqX | Scaffolds |
| *Ursus americanus* American Black Bear | ASM334442v1 | Male | 2588.39 | 100x | Illumina; PacBio | Scaffolds |
| Balaena mysticetus Bowhead Whale | * | | | 150 | Illumina HiSeq | Scaffolds |
| *Ursus arctos horribilis* Brown Bear | ASM358476v1 | Male | 2328.66 | 50x | Illumina HiSeq | Scaffolds |
| *Pan troglodytes* Chimpanzee | Clint_PTRv2 | Male | 3050.4 | 124x | Illumina HiSeq | Chromosomes |
| *Tursiops truncatus* Common Bottlenose Dolphin | Ttru_1.4 | Female | 2477.89 | 2.5x | Sanger; 454 FLX; Illumina HighSeq | Scaffolds |
| *Bos taurus* Cattle | ARS-UCD1.2 | Female | 2715.85 | 80x | PacBio; Illumina NextSeq 500; Illumina HiSeq; Illumina GAII | Chromosomes |
| *Canis lupus familiaris* Dog | CanFam3.1 | Female | 2407.29 | 7x | Sanger | Chromosomes |
| *Equus asinus asinus* Donkey | ASM30337v1 | Male | 2356.05 | 61x | Illumina | Scaffolds |
| *Loxodonta africana* African Savanna Elephant | Loxafr3.0 | Female | 3196.74 Mb | 7x | Sanger | Scaffolds |
| *Neophocaena asiaeorientalis* Yangtze Finless Porpoise | Neophocaena_asiaeorientalis_V1 | Male | 2284.63 | 106x | Illumina HiSeq2000 | Scaffolds |
| *Giraffa tippelskirchi* Giraffe | ASM165123v1 | Female | 2705.07 | 37x | Illumina HiSeq | Scaffolds |
| *Gorilla gorilla gorilla* Western Lowland Gorilla | GorGor4 | Female | 3063.36 | 80x | Illumina HiSeq | Chromosomes |
| *Eschrichtius robustus* Grey Whale | ASM218922v1 | Male | 2849.45 | 11x | Illumina HiSeq | Scaffolds |
| *Cavia porcellus* Domestic Guinea Pig | Cavpor3.0 | Female | 2849.45 | 6.8x | Sanger | Scaffolds |
| *Phocoena phocoena* Harbor Porpoise | ASM307100v1 | Female | 2571.07 | 87x | Illumina NextSeq 500 | Scaffolds |

| | | | | | | |
|---|---|---|---|---|---|---|
| *Monachus schauinslandi* Hawaiian Monk Seal | ASM220157v1 | Male | 2400.93 | 61x | Illumina HiSeq | Chromosome X and scaffolds |
| *Hippopotamus amphibius* Hippopotamus | ASM299558v1 | N/A | 2579.62 | 55x | HiSeq200 | Scaffolds |
| *Equus caballus* Horse | EquCab3.0 | Female | 2474.93 | 88x | Sanger; Illumina HiSeq; PacBio | Chromosomes |
| *Homo sapiens* Human | GRCh38.p13 | Both | 2987.97 | * | Sanger | Chromosomes |
| *Megaptera novaeangliae* Humpback Whale | megNov1 | Female | 2265.79 | 102x | Illumina HiSeq | Scaffolds |
| *Bos indicus x Bos taurus* Hybrid Cattle | UOA_Angus_1 | Male | 2630.86 | 136x | PacBio RSII; PacBio Sequel; Illumina | Scaffolds |
| *Tursiops aduncus* Indo-Pacific Bottlenose Dolphin | ASM322739v1 | Female | 2503.93 | 180x | Illumina HiSeq | Scaffolds |
| *Sousa chinensis* Indo-Pacific Humpbacked Dolphin | S_chinensis_fine_genome_map | Female | 2338.99 | 107.6x | Illumina HiSeq | Scaffolds |
| *Pteropus vampyrus* Large Flying Fox | Pvam_2.0 | Female | 2198.28 | 188x | Illumina | Scaffolds |
| *Myotis lucifugus* Little Brown Bat | Myoluc2.0 | Female | 2034.58 | 7x | Sanger | Scaffolds |
| *Trichechus manatus latirostris* Florida Manatee | TriManLat1.0 | Female | 3103.81 | 150x | Illumina HiSeq | Scaffolds |
| *Balaenoptera acutorostrata* Minke Whale | BalAcu1.0 | Male | 2431.69 | 92x | Illumina HiSeq 2000 | Scaffolds |
| *Mus musculus* House Mouse | GRCm38.p6 | N/A | 2689.66 | ** | Sanger | Chromosomes |
| *Heterocephalus glaber* Naked Mole-Rat | HetGla_female_1.0 | Female | 2631.08 | 90x | Illumina HiSeq | Scaffolds |
| *Monodon monoceros* Narwhal | NGI_Narwhal_1 | Male | 2414.06 | 42x | 10X Genomics; Dovetail Chicago; Dovetail | Scaffolds |
| *Okapia johnstoni* Okapi | ASM166083v1 | Male | 2878.13 | 30x | Illumina HiSeq | Scaffolds |
| *Monodelphis domestica* Gray Short-Tailed Opossum | MonDom5 | N/A | 3598.44 | 6.8x | Sanger | Scaffolds |
| *Orcinus orca* Killer Whale | Oorc_1.1 | Female | 2372.92 | 200x | Illumina HiSeq | Scaffolds |
| *Lagenorhynchus obliquidens* Pacific White-Sided Dolphin | ASM367639v1 | Female | 2334.47 | 35.68x | Illumina HiSeq | Scaffolds |
| *Ailuropoda melanoleuca* Giant Panda | ASM200744v1 | Female | 2363.89 | 70x | Illumina HiSeq | Scaffolds |
| *Sus scrofa* Pig | Sscrofa11.1 | Female | 2459.03 | 65x | PacBio | Scaffolds |
| *Ursus maritimus* Polar Bear | UrsMar_1.0 | Male | 2301.38 | 101x | Illumina Genome Analyzer II | Scaffolds |
| *Equus przewalskii* Przewalski's Horse | Burgud | Male | 2395.95 | 85.63 | Illumina HiSeq | Scaffolds |
| *Rattus norvegicus* Norway Rat | Rnor_6.0 | Both | 2743.3 | 3x | Sanger; SOLiD; PacBio | Chromosomes |

| | | | | | | |
|---|---|---|---|---|---|---|
| *Macaca mulatta* Rhesus Monkey | rheMacS_1.0 | Male | 2971.33 | 100x | PacBio Sequel | Chromosomes |
| *Procavia capensis* Cape Rock Hyrax | Pcap_2.0 | Female | 3749.9 | 107x | Illumina; Sanger dideoxy sequencing | Scaffolds |
| *Mesoplodon bidens* Sowerby's Beaked Whale | MesBid_v1_BIUU | N/A | 2797.69 | 32.4x | Illumina HiSeq | Scaffolds |
| *Physeter catodon* Sperm Whale | ASM283717v2 | Female | 2512.14 | 248x | BGISEQ-500 | Chromosomes and scaffolds |
| *Panthera tigris* Amur Tiger | PanTig1.0 | Male | 2391.08 | 99x | Illumina HiSeq 2000 | Scaffolds |
| *Choloepus hoffmanni* Hoffmann's Two-Fingered Sloth | C_hoffmanni_2.0.1 | Female | 3286.01 | 65x | Illumina | Scaffolds |
| *Odobenus rosmarus* Pacific Walrus | Oros_1.0 | Male | 2400.15 | 200x | Illumina | Scaffolds |
| *Bubalus bubalis* Water Buffalo | Bubbub1.0 | Male | 2836.17 | 119x | Illumina HiSeq 2000 | Scaffolds |
| *Leptonychotes weddellii* Weddell Seal | LepWed1.0 | Female | 3156.9 | 82x | Illumina HiSeq | Scaffolds |
| *Ceratotherium simum* Southern White Rhinoceros | CerSimSim1.0 | Female | 2666.62 | 91x | Illumina HiSeq | Scaffolds |
| *Camelus ferus* Wild Bactrian Camel | CB1 | Male | 2009.19 | 30x | Illumina GAIIx; 454 GS-FLX Titanium; SOLid 3 | Scaffolds |
| *Bos mutus* Wild Yak | BosGru_v2.0 | Female | 2703.27 | 130x | Illumina HiSeq; Illumina GA | Scaffolds |
| *Lipotes vexillifer* Yangtze River Dolphin | Lipotes_vexillifer_v1 | N/A | 2429.21 | 115x | Illumina HiSeq 2000 | Scaffolds |
| *Bos indicus* Zebu cattle | ASM293397v1 | Male | 2707.15 | 100x | 454; IonTorrent; Illumina NextSeq; Illumina MiSeq | Chromosomes |

- * not on NCBI
- ** genome coverage not available on NCBI
- Neophocaena Phocaenoides, NCBI txid34892 is no longer on NCBI
- Monachus Monachus, NCBI txif248254 is no longer on NCBI
